# Evidence of the ability of endocrine disrupting compounds to induce testicular germ cell cancer in humans

**DOI:** 10.1101/2023.12.22.573063

**Authors:** Nour Nicolas, Delphine Moison, Amandine Jampy, Quentin Masson, Nathalie Dechamps, Sébastien Messiaen, Sonia Abdallah, Stéphanie Pozzi-Gaudin, Alexandra Benachi, Damien Ulveling, Claire Francastel, Virginie Rouiller-Fabre, Gabriel Livera, Marie-Justine Guerquin

**Affiliations:** Université de Paris and Université Paris-Saclay, Laboratory of Development of the Gonads, IRCM/IBFJ CEA, UMR Genetic Stability Stem Cells and Radiation, Fontenay aux Roses, France; Université de Paris and Université Paris-Saclay,IRCM/IBFJ CEA, UMR Genetic Stability Stem Cells and Radiation, Fontenay aux Roses, France; Service de Gynécologie-Obstétrique et Médecine de la Reproduction, Hôpital A. Béclère, Université Paris Sud, Clamart, France; UMR7216 Épigénétique et Destin Cellulaire, CNRS, Université Paris Cité, F-75013 Paris, France

**Keywords:** *Testicular cancer*, Environmental endocrine disruptor, Germ cell, Male reproductive health, Human testis, DEHP, BPA

## Abstract

Testicular cancer is an increasing burden in modern societies and the most common malignancy among young adult males. Environment contaminants, especially endocrine disrupting compounds (EDC), may play a significant role in the development of these cancers through epigenetic alterations occurring during fetal and neonatal development. As long-term studies in humans and suitable experimental models with the potential to develop testicular cancer are lacking, no causal link can be established between endocrine disruptor exposure and testicular cancer incidence. Therefore, we developed an experimental model that recapitulates the differentiation of germ cells from primordial germ cells (pluripotency) into spermatocytes (meiosis) by using xenografted human fetal testis combined with germ cell transplantation into adult testis compartments. Using this model, we demonstrate that long-term fetal exposure (until 12 weeks) to a mixture of Di-2-ethylhexylphthalate (DEHP) and Bisphenol A (BPA), two most prevalent plasticizers, could interfere with fetal germ cell differentiation, leading to carcinogenesis and seminomas. Transcriptome, methylome, and histological analyses reveal that BPA/DEHP exposure induced some significant hallmarks of germ cell tumors to occur: persistent pluripotent and proliferative germ cells, global hypomethylation of CpGs in germ cells, abnormal expression of meiotic markers and fibrotic signatures in fetal testis. Additionally, we found that EDC-exposed fetal germ cells were more likely to develop seminoma in a context that allows spermatogenesis to begin. This study proposes the first experimental evidence that EDC exposure can cause long-term, irreversible lesions in fetal germ cells, which then lead to testicular tumorigenesis in adults.

## INTRODUCTION

Testicular cancer incidences have steadily increased worldwide since 1990 and were correlated with socio-demographic factors (Tian et al., 2023). Testicular germ cell tumors (TGCTs) are the most common cancer in teenagers and young adults (15-40 years-old) (Rajpert-De Meyts et al., 2016). TGCTs are heterogeneous neoplasms arising from neoplastic transformation of germ cells with various degrees of differentiation. They are divided into carcinoma in situ (CIS, the preinvasive stage of testicular germ cell tumours), seminomas, and nonseminomas. Seminoma and CIS have close similarity to fetal primordial germ cells (PGCs), the precursors of gametes. First, the proliferative tumor cells express a variety of stem cell factors, including OCT3/4, NANOG, PLAP, and SOX2. Second, tumor cells present global DNA hypomethylation compared to other somatic-derived cancers (Hart et al., 2005; Hoei-Hansen et al., 2005; Jones et al., 2004). Hence, seminoma and CIS are believed to have embryonic origins and could result from PGC differentiation failure (Rajpert-De Meyts et al., 2016). Human PGC that emerge from the proximal epiblast migrate into the developing gonad at 5-6 weeks post-fertilization (WPF). Prior to integration into the somatic niche, PGCs are highly proliferative. They retain pluripotency, producing many stem cell factors, including OCT3/4 (encoded by POU5F1), NANOG, SOX2, AP2γ (TFAP2C) and PDPN. Due to global DNA demethylation in germ cells and somatic paracrine pressure, PGCs (from 9 WPF) down-regulate pluripotency factors and turn on DDX4, DND1, and meiotic genes as SYCP3 and DAZL that enable them to undergo sexual differentiation and gametogenesis (Hu et al., 2015; Li et al., 2021; Tang et al., 2016). During this stage and just after NANOS2 expression, differentiating PGCs gradually slow down their proliferation. They get closer to the basement membrane of testis cords, express MAGE-A4 illustrating their differentiation, and are called pre-spermatogonia. During the quiescent phase, DNA *de novo* methylation occurs. After birth, germ cells (GCs) resume proliferation and spermatogenesis. The asynchronous transition from PGCs expressing pluripotency factors to a more differentiated phenotype is a key event in fetal testis development and infertility, chromosomal abnormalities, and germ cell tumors can occur when disrupting this finely regulated transition.

The genesis of TGCTs could be driven by genetic factors, but environmental factors is also suspected to influence malignant transformation (Litchfield et al., 2017; Lobo et al., 2019). Due to the putative carcinogenesis and strong association with other reproductive disorders, environmental endocrine disrupting compounds (EDCs) have been blamed for the rise in testicular cancers (Ghazarian et al., 2018). It is difficult, however, to establish a causal association between EDCs and testicular cancer in humans due to a lack of long-term studies across a wide caseload. Only short term studies suggested that fetal germ cell differentiation could be impaired by EDC exposure. Among EDCs studied, bisphenols and phthalates were shown to impair PGC differentiation and induce germ cell apoptosis and cytokinesis impairment (Eladak et al., 2018; Li et al., 2022; Abdallah et al., 2023; Kilcoyne and Mitchell, 2019). The most widely used phthalate ester is di-(2-ethylhexyl) phthalate (DEHP), which is incorporated in plasticizers and other products (Li et al., 2022). Hard plastics are commonly manufactured with bisphenols, synthetic chemicals. Epoxy resins, which coated metallic food and beverage cans, also contain bisphenol A (BPA). Hence, in the manufacture of consumer and medical products, phthalates and bisphenols are high production volume chemicals. Due to the widespread presence of bisphenol and phthalates in the environment, 75-90% of the general population is commonly contaminated with these chemicals, according to biomonitoring studies (Li et al., 2022; Ramadan et al., 2020). As a result of their omnipresence (20 to 99% of cord blood samples contain BPA and DEHP), they are known to affect pregnancy rate, fetal development, and, ultimately, GC differentiation.

The purpose of this study was to establish a causal link between fetal exposure to EDCs and the incidence of TGCT. Due to the high prevalence and reprotoxicity of BPA and DEHP, it seems obvious to evaluate the incidence of chronic *in utero* exposure to a mixture of BPA and DEHP on TGCT formation. Human fetal testis tissue xenograft allowed testicular development and maintenance during a long period (Eladak et al., 2018; Kilcoyne and Mitchell, 2019). Using this model, we exposed human fetal testis *via* nude mice to a mixture of BPA and DEHP (DE/BP, 1µM each) for 7-10 weeks. Hallmarks of CIS/seminoma have been retrieved at the end of exposure after histological, transcriptomic and epigenomic (DNA methylation) analysis. Germ cell differentiation is impaired by DE/BP exposure, which maintains GCs at the proliferative stage. We also observed global DNA hypomethylation in PGCs but also in pre-spermatogonia and hypomethylated genes in pre-spermatogonia are linked to TGCTs. Impairment of peritubular myoid cells and fibrotic response were also induced by DE/BP exposure and could be linked to fetal GC differentiation impairment. Finally, heterologous transplantation of human treated fetal GC into adult murine testes has been performed to evaluate the tumorigenic potential of DE/BP treated germ cells. The previous DE/BP treatment appears to have affected the ability of human GC to homing in to murine somatic and spermatogenesis. Interestingly, neoplastic formation and seminoma rise was also observed after DE/BP-treated GC transplantation.

## METHODS

### Ethics statement

Fetal human fetal testes (8-35 weeks post-fertilization [WPF]) were obtained from pregnant women referred to the Department of Gynecology and Obstetrics at the Antoine Béclère hospital (Clamart, France) for legally induced abortions (first trimester, 8-11 WPF) or therapeutic termination of pregnancy (second-third trimester, 25-35 WPF) as previously described (Eladak et al., 2018; Fouquet et al., 2017). None of the abortions were due to a fetal abnormality. All women provided written informed consent for the scientific use of fetal tissues. The project was approved by the local Medical Ethics Committee and the French Biomedicine Agency (reference number PFS 12–002).

### Animals

Adult athymic mice (NMRI-Foxnl^nu^/foxnl^nu^, from Janvier labs) were used IN xenograft and germ cell transplantation experiments. Mice were housed in accordance with National Institutes of Health guidelines in our animal facility under controlled photoperiod conditions (lights on from 08:00 am to 08:00 pm). They were supplied with estrogen-free commercial food and tap water ad libitum. Mice received water via glass water bottles. The animal facility is licensed by the French Ministry of Agriculture (agreement n° B92-032-02). All animal care protocols and experiments were reviewed and approved by the ethics committee of CETEA–CEA DSV (France, APAFIS 19466-2019022610468495 v2) and followed the guidelines for the care and use of laboratory animals of the French Ministry of Agriculture. During surgical procedures, mice were anaesthetised by isoflurane inhalation, placed on a heating plate at 37 °C to avoid anaesthesia-induced hypothermia and constantly perfused with saline solution to prevent tissue deshydration. Mice were previously injected with buprenorfine (0.5 mg/kg) and the scar site was previously treated with a lidocaine cream. During one week post-surgery, mice received analgesia (Carprofen, 25 μg/mL) in the drinking water.

### Histology & Immunostaining

Tissues (grafted human or transplanted mouse testes) were fixed in Bouin’s fluid during 2 hours at room temperature, dehydrated, embedded in paraffin, and cut into 5 μm thick sections. After rehydration, testes section were subjected hematoxylin-eosin or Trichrome’s Masson staining (standard protocols) or immunostainings. Primary antibodies used in this study and their related retrieval and blocking protocols were reported in Supplementary Table 1. For immunofluorescence, slides were autoclaved with 10 mM sodium citrate buffer (pH 6.0) or 10 mM Tris/EDTA (pH 9.0) and blocked in 5% BSA with 0.1% Triton X-100, 2.5 % normal horse serum (Impress HRP Reagent Kit MP-7402) or 2% gelatin, 0.05% Tween and 0.2% BSA (gelatin buffer) for 1 h before adding antibody. Primary antibody was incubated at 4°C overnight and slides were washed 3 times in a washing buffer. Then, specific donkey secondary antibody conjugated with AlexaFluor (384,488, 594 nm at 1:500) was applied for 1 hour at room temperature. Slides were counterstained with DAPI (1/20000) and mounted in ProlonGold. A Leica TCS SP8 (confocal) or Leica DM5500 B (epifluorescence) were used for fluorescence observation/acquisition. Images and methylation fluorescence intensity were analyzed with Image J software. For immunochemistry, slides were autoclaved with 10 mM sodium citrate buffer (pH 6.0) or 10 mM Tris/EDTA (pH 9.0). After inactivation of endogeneous peroxidase activity with 0.3% H2O2 in water (room temperature, 30 min), slides were subjected to blocking solution (2.5 % NHS, 30min at room temperature). A specific secondary HRP-antibody (Impress, Vector laboratories) was incubated during 30 min at room temperature. HRP activity was revealed with DAB and counterstained with haematoxylin. Slides were dehydrated and mounted with Eukit. Olympus BX51 was used for image acquisition coupled with Histolab software.

### Xenografting procedures and EDC treatment

Xenografting procedure was performed as previously described (Eladak et al., 2018). Briefly, human fetal testis (from 8-11 WPF) were cut into small pieces (∼1 mm^3^) and 4-6 pieces/testis (depending of the age or size of the gonad) were insert into the muscle of the left back of two twin female mice. One week after surgery, one mouse was exposed by drinking water to a mixture of BPA (1 µM, CAS no. 80-09-1, Merck) and DEHP (1 µM, CAS no. 117-81-7, Merck) and the other was exposed by drinking water to the vehicle (absolute ethanol diluted at 1/10^6^ in water). Daily intake of DEHP and BPA was evaluated to ∼35 µg/kg and ∼58 µg/kg respectively. Mice were exposed during 6-11 weeks (corresponding to the 18 WPF equivalent age) and at the end of exposure, testes were collected and each pieces were cleaned from extracellular matrix and murine muscular tissue. One part of the pieces were used for histological studies, one the other part for transcriptome and methylome analyses and stored at-80°C in RLT plus until DNA/RNA extraction and the last part of testis pieces were used for germ cell transplantation.

### Human cell sorting and transplantation

After xenograft collection, tissue was gently dissociated using 21g needles in accutase solution (A6964, Merck) and incubated at 37°C during cell dissociation. Twenty minutes later, fetal calf serum (10 %, Merck) was added and cells were centrifuged for 10 min at 800 g. Cells were resuspended in labelling solution containing anti-human FITC Beta2 microglobuline (1/10; 551338, BD biosciences) diluted in PBS-BSA 0.5 % and incubated for 30 min at room temperature. Cell suspension was filtered (20 µm) and Hoechst 33358 (cell viability) was added before cell sorting with BD Influx sorter. B2M negative cells were resuspended in PBS supplemented with 2.5 mg/ml DNase I (DN25, Merck) and 10 % trypan blue solution (T8154, Sigma-Aldrich) for transplantation.

To deplete endogenous spermatogenesis, 4-week-old recipient male nude mice received an intraperitoneal injection of busulfan (40 mg/kg). At least four weeks later, mice were used for human germ cell transplantation. Both testes and epididymes were exteriorized through a midline incision in their abdomen and 10 µl solution of donor B2M negative cells (containing 100 000 cells) was micro-injected *via* efferent duct into the seminiferous tubules of the testis of the recipient mouse as previously described (Medrano et al., 2014). One mouse is injected with DE/BP-exposed cells and the other with vehicle-exposed cells. In the absence of the observation of apparent extratesticular tumor (subcutaneous mass or skin lesion, Supplementary Figure 9, recipient mice were euthanized four weeks after transplantation and testis were collected for histological analyses.

### B2M cells RNA isolation and quantitative PCR

Total RNA was extracted from B2M cells using the RNeasy kit (QIAGEN), reverse-transcribed using High Capacity Reverse Transcription Kit (Thermofisher Scientific), and processed for qRT-PCR with PowerUp SYBR Green Master Mix (Applied Biosystems) and a StepOne system (Thermofisher Scientific) with gene-specific probes (B2M: 5’-CACAGCCCAAGATAGTTAAGT & 5’-CCAGCCCTCCTAGAGC, the other probes have been already described in (Arkoun et al., 2022; Frydman et al., 2017; Le Bouffant et al., 2010). Reactions were run in triplicate and mRNA levels were normalized to ACTB1 and quantified using the delta-delta Ct method. The values shown are mean ± s.e.m. from three biological replicates.

### Xenograft DNA/RNA isolation and quantification

Total DNA and RNA from xenografted testes were simultaneously isolated using the AllPrep DNA/RNA microkit (ref: 80284, Qiagen, France) according to manufacturer’s instructions. DNA and RNA concentrations and quality were assayed with Qbit fluorometer (Thermofisher) and Agilent bioanalyser. After DNA and RNA quantitation, pools of DNA or RNA per condition were performed containing from 3 to 8 testis per pool.

### RNA sequencing, alignment and differential analysis

RNA sequencing was carried out in triplicate under condition (CTL and DE/BP). RNA-Seq libraries are prepared by Integragen SA (Evry, France) with NEBNext® Ultra™ II Directional RNA Library Prep Kit for Illumina according to supplier recommendations (NEB). The capture is then performed on cDNA libraries with the Twist Human Core Exome +Custom IntegraGen Enrichment System according to supplier recommendations (Twist Bioscience). Then a capture of the transcriptome coding regions is performed and the resulting library is suitable for subsequent cluster generation and sequencing. Briefly, the RNA is fragmented into small pieces using divalent cations under elevated temperatures. CDNA is generated from the cleaved RNA fragments with random priming during first and second strand synthesis and sequencing adapters are ligated to the resulting double-stranded cDNA fragments and enriched by 7 PCR cycles. The transcriptome coding regions are then captured from this library using sequence-specific probes to create the final library. For that purpose 500 ng of purified libraries are hybridized to the Twist oligo probe capture library for 16 hr in a singleplex reaction. After hybridization, washing, and elution, the eluted fraction is PCR-amplified with 8 cycles, purified and quantified by QPCR to obtain sufficient DNA templates for downstream applications. Each eluted-enriched DNA sample is then sequenced on an Illumina HiSeq4000 as 75b reads. Image analysis and base calling is performed with Illumina Real Time Analysis (2.7.7) with default parameters. Raw sequencing data was quality-controlled with the FastQC program. Low-quality reads were trimmed or removed using Trimmomatic (minimum length: 50 bp). Paired-end-reads were aligned to the human reference genome (hg38 build) with STAR software. Mapping results were quality-checked using RNA-SeQC. Gene counts were obtained using rsem tools (rsem-calculate-expression, option for paired-end and stranded). Normalization and differential analysis were performed with the edgeR package. Functional Enrichment analysis was carried out using EnrichR and Cluster Profiler packages.

### RRBS, alignment, methylation calling and differential analysis

RRBS sequencing was performed in triplicate by condition (CTL and DE/BP). RRBS was performed by Integragen SA (Evry, France) with the Diagenode Premium RRBS kit. In brief, 100 ng of qualified genomic DNA were digested with MspI. After end-repair, A-tailing, and ligation to methylated and indexed adapters, the size selected library fragments were subjected to bisulfate conversion, PCR amplified, and sequenced on an Illumina HiSeq4000 sequencer as Paired_End 75 bp reads. Image analysis and base calling is performed using Illumina Real Time Analysis (2.7.7) with default parameters. Base calling is carried out with using the Real-Time Analysis software sequence pipeline (2.7.7) with default parameters. BS-Seeker2 (v2.1.8) was employed to map RRBS data to the human genome (hg38) and retrieve the number of methylated and unmethylated cytosines at each covered CpG site. Allowing local/gapped alignment with Bowtie2, increased mappability. The parameters used are: -r (Map reads to the Reduced Representation genome), -c C-CGG (MspI: sites of restriction enzyme and specifying lengths of fragments ranging [40bp, 400bp]. One mismatch is allowed in the adaptor sequence. The module bs_seeker2-call_methylation.py was used to call methylation levels from the mapping result with these parameters: --rm-SX (Removed reads which would be considered as not fully converted by bisulfite) and –rm-overlap (Removed one mate if two mates are overlapped, for paired-end data).

### Single Cell analysis for bulk RNAseq and RRBS data integration

Published single cell datasets from the same study (Garcia-Alonso et al., 2022) of human fetal ovary (33 datasets) and testis (22 datasets) from 6 to 21 WPF were used for this analysis. Cell-by-gene matrices for each sample were individually imported to Seurat version 4.0.4 and aggregated by sex. Male germ and somatic cells population were extracted from male merged object using a expression cut-off of genes considered as germ and somatic cell marker (male GC: DDX4, POU5F1, DDX4, DAZL, MAGEA4, SYCP3, STRA8 or NANOS2, supporting cells: SOX9, AMH, PAX8, MYH11, GATA4, INSL3, ARX, NR5A1, Lymphoid/myeloid/endothelial cells: HBA1, CDH5, PTPRC). Female germ cells were extracted from female merged object using the same cut-off of male germ cells. Cells with an unusually high or low number of UMIs or mitochondrial gene percent (>10%) are excluded. All male germ cells and only 400 supporting/immune cells were used in this study). All female/male germ cells and somatic male germ cells were merged for further clustering. For single cell gene signature we calculate the module score using a list of up- or down-regulated genes (abs(LFC) ≥ 0.5 and FDR ≤ 0.05). Determination of the expression thresholds (“high”, “medium” and “low” levels) of up- and down-regulated genesets suitable for bRNAseq and RRBS integration was determined based of the distribution of cell density presented in Supplementary Figure 4B.

## RESULTS

### DE/BP exposure delays fetal germ cell differentiation in the human fetal testis

To mimic environmental EDC *in utero* exposure, we examined the adverse effects of long-term exposure to a mixture of EDCs on fetal testicular human germ cells. Two female nude mice were xenografted with human fetal testes (from 8-11 WPF) and exposed to DEHP and BPA (DE/BP, 1 m each) or vehicle (CTL, ethanol at 1/106) by drinking water for 6-11 weeks, depending on their embryonic stage when collected. At equivalent stage of 18 WPF, xenografted testes were collected and germ cell survival and differentiation were evaluated (Figure 1, Supplementary Figure 1). During the second half of pregnancy, *in vivo* human fetal germ cell differentiation is characterized by a gradual decrease of pluripotent proliferative primordial germ cells expressing OCT-3/4, AP2γ, KIT or PDPN and an increase of DDX4- and quiescent pre-spermatogonia as shown in Supplementary Figure 1. Testicular xenografting did not prevent germ cell differentiation as we observed an increase of DDX4- and MAGEA4-positive germ cells and decrease of POU5F1- and AP2γ-positive germ cells in DE/BP and CTL treated testes compared to 8-11 WPF ungrafted testes (Figure 1A,B). However, DE/BP exposure delayed germ cell differentiation as we observed a significant increase in OCT3/4-, AP2γ-, KIT- and PDPN- positive germ cells and decrease of DDX4-positive germ cells in DE/BP-treated testes compared to CTL-treated testes (Figure 1A-B, Supplementary Figure 2). No significant change in MAGE-A4 germ cell proportion was observed in DE/BP testis (Figure 1A). To understand the modification of the ratio of undifferentiated/differentiated germ cells in DE/BP exposure, we evaluate germ cell survival and apoptosis. DE/BP testis had similar germ cell numbers to CTL testis, and apoptotic germ cells (cleaved caspase 3 positive germ cells) were not increased after DE/BP exposure (Figure 1C-D).

**Figure 1.**
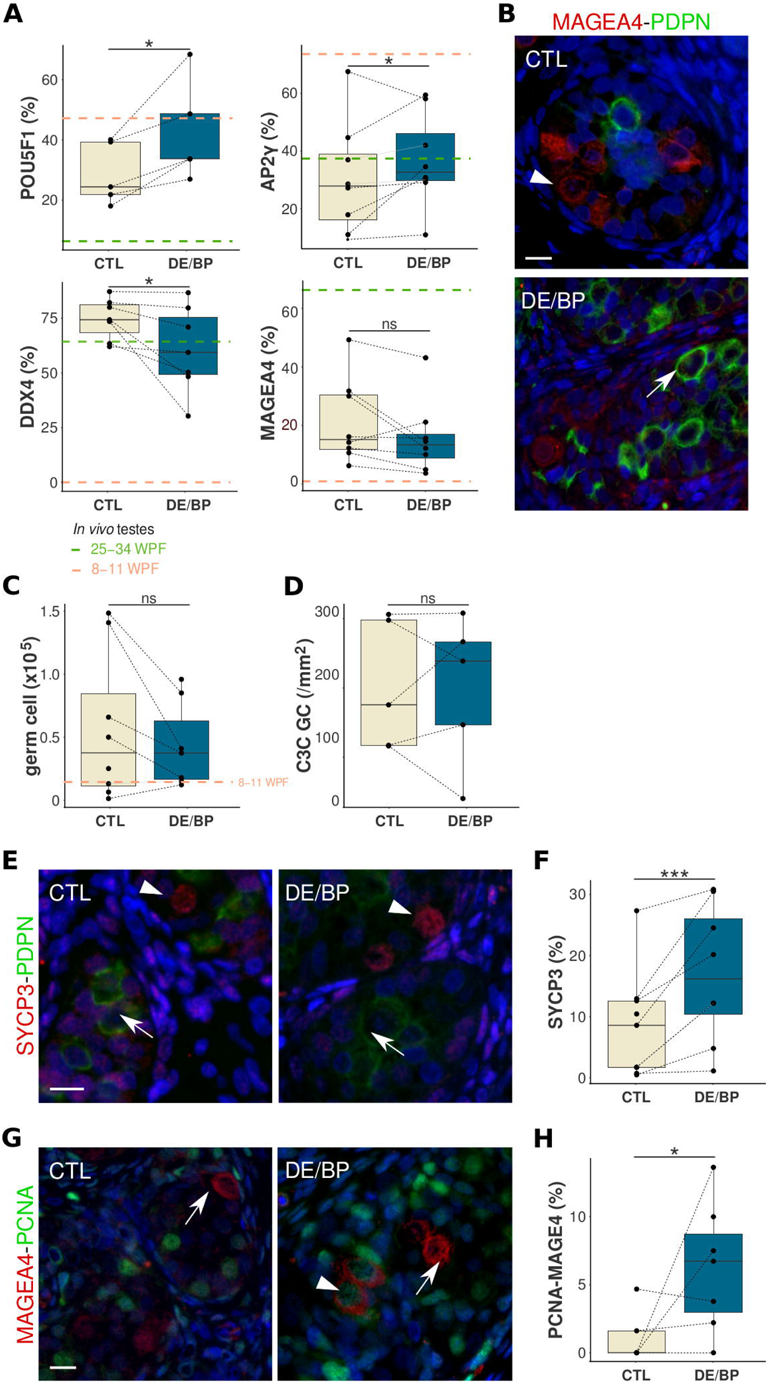
Germ cell survival and differentiation after DE/BP exposure. **(A)** Percentage of POU5F1 (OCT-3/4), AP2γ, DDX4 and MAGEA4 positive GCs (/total GCs) from control (CTL) and DEHP-BPA (DE/BP) xenografted testes. **(B)** Representative images of xenografted testes stained with anti -PDPN (green) and -MAGEA4 (red) antibodies counterstained with DAPI (blue). Arrow indicates PDPN positive GC and arrowhead indicates MAGEA4 positive GC. **(C)** Germ cell count from CTL and DE/BP xenografted testes. **(D)** Density of Cleaved 3-Caspase (C3C) positive germ cells (GC, AMH negative cells in testis cords) from CTL and DE/BP xenografted testes. **(E)** Representative images of testes stained with anti -PDPN (green) and -SYCP3 (red) antibodies counterstained with DAPI (blue). Arrow indicates PDPN positive GC and arrowhead indicates SYCP3 positive GC. **(F)** Percentage of SYCP3 positive GCs (/total GCs) from CTL and DE/BP xenografted testes. **(G)** Representative images of testes stained with anti -PCNA (green) and -MAGEA4 (red) antibodies counterstained with DAPI (blue). Arrow indicates non-proliferative PCNA negative – MAGEA4 positive GC and arrowhead indicates proliferative PCNA positive – MAGEA4 positive GC. **(H**) Percentage of PCNA-MAGEA4 postive GCs (/total MAGEA4 positive GCs) from CTL and DE/BP xenografted testes. (A-B,D, F & H) box plot (median and interquartile ranges) from 6-8 paired xenografted testes. * p ≤ 0.05, ns: non significant (Paired student’s t test). Grey dotted line connects paired xenografted testes (indicated by points). Orange dotted lines and green dotted lines represent mean of count or percentage from 8-11 WPF and 25-34 WPF respectively *in vivo* testes. (C, E, G). Bar represents 10 µm.

*In vivo,* germ cell differentiation toward the male pathways (i.e., from gonocyte to prespermatogonia) is characterized by the repression of the meiotic program, MAGEA-4 expression, and cell cycle arrest thereafter. We observed a significant increase in SYCP3-positive germ cells in DE/BP testes, indicating that some male germ cells can escape meiosis (Figure 1E-F). Furthermore, DE/BP testes had significantly more proliferative MAGEA4-positive germ cells (Figure 1G-H) and this abnormal maintenance of proliferation in MAGEA-4 germ cells could explain the nonsignificant difference in MAGEA-4 expressed cells between CTL and DE/BP testis. Altogether, the findings suggest that long-term exposure to a mixture of DEHP and BPA delays male germ cell differentiation and leads to some defects in germ cell differentiation, as evidenced by the production of meiotic markers in male germ cells and proliferation maintenance in pre-spermatogonia.

### DE/BP exposure induces transcriptional modification in the human fetal testis

In order to identify transcriptional responses to DE/BP exposure in fetal testes, bulk RNA sequencing (bRNAseq) was performed on DE/BP- and CTL-xenografted testes (Figure 2). Because bulk RNA-seq averages out gene expression across all cells in the sample, we also assessed the testicular cell response to DE/BP using single cell RNA sequencing (scRNAseq, Supplementary Figure 3). For unknown reasons, low cell counts and high ambient RNA were observed in DE/BP testes after cell encapsulation. Comparing DE/BP testes with CTL testes, we observed a 90 % loss of total cells. Therefore, the number of GCs and some somatic cells was insufficient to accurately assess their transcriptional response to DE/BP (Supplementary Figure 3A-B). We focus on bRNAseq analysis in our study. The differential expression analysis reveals a global down-regulation of genes in DE/BP-treated testis compared to CTL testis (2561 genes with log fold change -0.5 and *AdjpVal* ≤ 0.05, Figure 2A). An analysis of functional enrichment of down-regulated gene set revealed a strong enrichment of genes associated with *DNA Repair* (GO:0006281, *AdjpVal* = 2.44 E-08, Supplementary Table 2) and *Double-Strand Break Repair Via Homologous Recombination* (GO:0000724, *AdjpVal* = 1.39 E-05, Supplementary Table 2), which may be involved in meiosis and germ cell development. Using published single-cell transcriptomes from human fetal testis and ovary, we projected on down- and up-regulated genesets sorted by differential analysis to clarify which testicular cell types are more affected by EDCs exposure. Single cell datasets (cell-by-gene matrices, (Garcia-Alonso et al., 2022) from human fetal ovary and testis (from 6 to 21 WPF) were used to cover somatic and germ cell fetal differentiation. Total male and female germ cells were extracted and added to male somatic cells to perform UMAP projection and supervised clustering (Figure 2B and Supplementary Figure 4A). For germ cells, we differentiated four clusters: undifferentiated and pluripotent male and female germ cells (expressing pluripotent genes as *POU5F1*), differentiating male and female primordial germ cells (expressing *DDX4*, *DAZL* and *STRA8*), meiotic oocytes (from ovary only, highly expressed meiotic genes as *SYCP3*) and quiescent pre-spermatogonia (from testis only, expressing *MAGE-A4* and *NANOS2*). For testicular somatic cells, we differentiated 3 clusters: supporting, interstitial and “immune” cells. Supporting cells included immature (*SOX9-* positive cells), mature sertoli cells (*AMH-* positive cells) and *PAX8-*positive cells from the *rete testis.* Interstitial cells included myoid/peritubular (*SMA-* and *COL6A1-* positive cells) and leydig cells (*NR2F2*- and *STAR-*positive cells). “Immune” cells included HBA1-positive myeloid cells, PTPRC-positive lymphoid cells and CDH5-positive endothelial cells. Thanks to single cell transcriptome data, we identified that down-regulated genes observed in DE/BP testes were preferentially highly expressed by germ cells, mostly in differentiating PGCs and quiescent pre-spermatogonia (Figure 2B-C). On the contrary, up-regulated genes were mosly expressed by somatic cells, especially in interstitial cells (Figure 2B-C). The decrease of transcripts quantified by bRNAseq normally expressed in differentiating PGCs and pre-spermatogonia in DE/BP-treated testis seems to result of differentiated germ cell pool decrease as previously evidenced in Figure 1. Indeed, identified markers differentially expressed in quiescent pre-spermatogonia or differentiated PGCs (Log fold chance ≥ 0.5, *AdjpVal* ≤ 0.05 in comparison to all others clusters) were mostly under-represented in DE/BP testes compared to CTL (22- 23 % of all markers of pre-spermatogonia and differentiating PGCs, Figure 2D). Functional enrichment analysis of down-regulated genes in DE/BP testes and differentially expressed (compared to others somatic cell clusters) in all GC cluster (*ie “*undifferentiated PGC”, “differentiating PGC” and “meiotic oocyte” and “quiescent pre-spermatogonia”) showed a strong enrichment of genes involved in meiosis initiation and meiotic homologous recombination such as *SYCP3, SYCP1*, *MEIOB or MEIOC* (Figure 2E, Supplementary Figure 4C and Supplementary Table 3). During the differentiation process called meiotic licensing, PGC switch-off their pluripotency program and adopt a transcriptional program allowing post-natal gametogenesis and, by extension, meiosis. During this switch, male differentiating PGCs over-expressed meiotic genes, but male somatic environment prevents the activation of meiosis initiation until post-natal life. For this reason, we suggest that down-regulated genes retrieved from pathways presented in Figure 2E and normally expressed during PGC differentiation (Figure 2F) correspond to an arrest or delay of germ cell differentiation in DE/BP treated testis. However, functional analysis of up-regulated DEGs specifically expressed by somatic cells (*ie* “supporting cells”, “interstitial cells” and “immune cells”, Figure 2G) suggested an up-regulation of genes associated with extracellular matrix, fibrillogenesis and fibrosis. Differential analysis using scRNAseq data on supporting and interstitial cells confirmed the up-regulation of genes involved in ECM production, focal adhesion and TGF-β signaling (Supplementary Figure 3E). In humans, peritubular myoid cells (PMCs) form several thin layers (5-8 layers) surrounding testis cords. These cells also secrete components of the extracellular matrix of the peritubular wall as collagens and proteoglycans. Accumulation of ECM proteins and PMC hypertophy are hallmarks of tubular fibrosis and are correlated to spermatogenesis defects and testicular cancer (Mayerhofer, 2013). After DE/BP exposure, we did not evidence any significant increase of ECM deposit but PMC proliferation and number of PMC layers seems to be affected ( Supplementary Figure 5 A-C). Altogether, these results confirm that DE/BP exposure interfere with PGC differentiation transiently maintaining germ cells at the pluripotent stage. Moreover, our transcriptome analysis reveals that somatic cells, and particularly PMCs or precursors of PMCs, were subjected to stress leading to TGF-β dependent fibrotic response.

**Figure 2.**
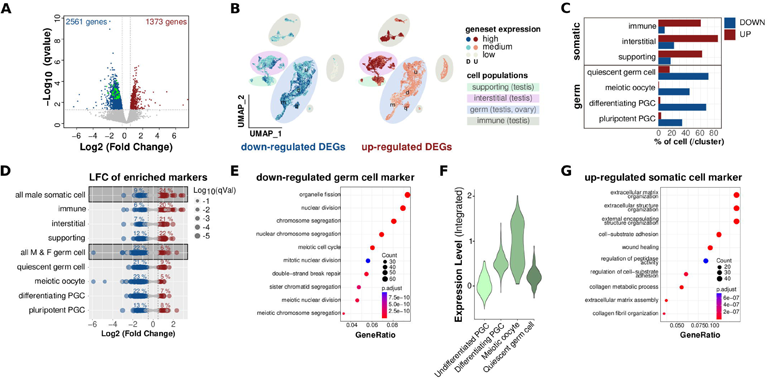
Bulk RNAseq analysis illustrates the delay of GC differentiation and reveals somatic response leading to extracellular matrix modifications. **(A)** Volcano plot of bulk RNA-seq analysis of differently expressed genes from DEHP/BPA (DE/BP) vs control (CTL) xenografted whole testes. Red dots indicate up-regulated genes (LFC ≥ 0.5, qValue ≤ 0.05). Blue dots indicate down-regulated genes (LFC ≤ −0.5, qValue ≤ 0.05). Green dots indicate genes involved in *Double-Strand Break Repair Via Homologous Recombination (GO:0000724).* (B) Uniform manifold approximation and projection (UMAP) clustering of combined testicular (somatic and germ) and ovarian (germ only) cells from fetal human gonads presenting the expression level (High, medium and low) of downregulated (blue) and upregulated (red) genesets. As presented in Supplementary Figure 4, supporting cluster (2049 cells) included AMH positive sertoli cells, SOX9 positive sertoli cells and PAX8 positive cells from rete testis. Interstitial cluster (3259 cells) included NR2F2, ARX positive interstitial progenitors, SMA, LUM, COL6A1 positive myoid cells, INSL3/STAR positive leydig cells. Immune cluster (3007 cells) included HBA1 positive myeloid cells, PTPRC positive lymphoid cells and CDH5 positive endothelial cells. Germ cluster included POU5F1 (OCT3/4) positive pluripotent male and female germ cells (u, 2908 cells), DDX4/STRA8 positive, MAGEA4 negative differentiating male and female germ cells (d, 2674 cells), SYCP3 positive meiotic oocyte (m, 3168 cells) and MAGEA4/NANOS2 positive quiescent pre-spermatogonia (“quiescent germ cell”, q, 404 cells). **(C)** Proportion of cell by cluster expressing high level of up- or down-regulated genesets after DE/BP exposure observed in bulk RNA-seq analysis. **(D)** Dot plot showing Log2 Fold-change (LFC) values from RNA-seq analysis (DE/BP vs CTL) of gene markers of cell clusters (differentially up-regulated) obtained from sc-RNAseq analysis. Size of dots correspond to the qvalue obtained from bulk RNA-seq differential analysis. **(E)** Functional enrichment analysis of down-regulated genes after DE/BP exposure and specifically expressed in germ cell population (u, d, m and q populations). **(F)** Mean expression of genesets retrieved in enriched pathways observed in E (from “organelle fission” to “meiotic chromosome segregation”). **(G)** Functional enrichment analysis of up-regulated genes after DE/BP exposure and specifically expressed in supporting, interstitial and immune clusters.

### DNA methylation modification induced by DE/BP exposure in the fetal human testis

Epigenomic studies on TGCT revealed that global DNA methylation is low in seminomas and resembles to DNA methylation pattern observed in PGC (Shen et al., 2018). For this reason, we evaluated DNA methylation in DE/BP and CTL xenografted testes (Figure 3). DNA demethylation-remethylation waves occurs during fetal germ cell differentiation (Guo et al., 2017). We first assayed DNA methylation status by immunodetection of 5-methylcytosine (5mC) and 5-hydroxymethylcytosine (5hmC, an intermediate metabolite during DNA methylation) considering the differentiation state of GCs (Figure 3A and Supplementary Figure 6). As pre-spermatogonia differentiation is concomitant with migration to the basal compartment (Supplementary Figure 6A), we have categorized GCs depending on their localization in testis cords considering central GCs as PGCs and peripheral GCs as pre-spermatogonia. Analysis of fluorescence intensity of 5mC and 5hmC in somatic and germ cells showed that global DNA methylation intensity (5mC plus 5hmC fluorescence intensity) was significantly decreased in basal and central GCs after DE/BP exposure (Figure 3A and Supplementary Figure 6B-C). No significant difference of DNA methylation intensity between DE/BP and CTL testes was observed in somatic cells ( Supplementary Figure 6B). ITo characterize the DNA methylation status at a genome-wide level, we performed Reduced Representation Bisulfite Sequencing (RRBS) of DE/BP and CTL xenografted bulk testes at the end of exposure. Differential methylation analysis showed that genomic regions were mostly differentially hypomethylated in DE/BP testis compared to CTL (932 DMRs hypomethylated VS 128 DMRs hypermethylated, Figure 3B). At single loci levels, we also observed that CpGs were preferentially differentially hypomethylated in DE/BP testes compared to CTL (18 817 CpGs hypomethylated vs 9297 CpGs hypermethylated, data not shown). Genes that showed more than 50 % of differentially hypomethylated CpGs were most abundant than genes with more than 50 % of differentially hypermethylated CpGs (7.0 % vs 4.3 % of genes, Figure 3C). CpG hypomethylation observed in DE/BP treated cells affects all gene regions (promoter, exon, intron and 3’-end, Figure 3C). As previously mentioned for bRNAseq, bulk RRBS average methylation of all cells. For this reason, we also integrated differential analysis methylome data to scRNAseq to determine witch cell population was more damaged by gene hypomethylation. We observed that genes with CpGs mostly hypomethylated were preferentially expressed by quiescent pre-spermatogonia suggesting that these cells will be more affected by epigenomic modification by DE/BP exposure (Figure 3C). An interesting findings was the functional association of GC hypomethylated genes (from pluripotent GCs to quiescent spermatogonia) with rare disease was linked to cancers as *testicular cancer* and *retinoblastoma* (Figure 3E). Low correlation between methylation status and transcriptional regulation was observed but genes with significant hypomethylation in the promoter region and transcriptional up-regulated could be linked to carcinoma disease (Supplementary Table 3-4). Overall, DE/BP exposure led to global hypomethylation, particularly in differentiating PGCs and pre-spermatogonia leading to transcriptional regulation and carcinogenesis induction.

**Figure 3.**
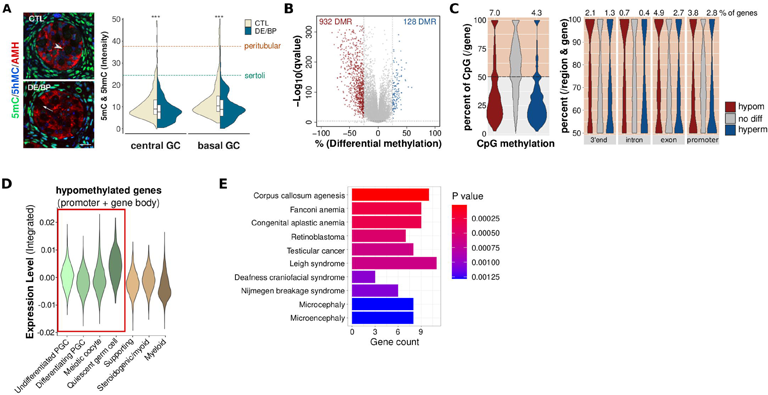
DNA methylation changes in germ cell after DE/BP exposure. **(A)** Left: Representative images of testes stained with anti −5mC (green), −5hmC (blue) and -AMH (red) antibodies. Arrowhead indicates central germ cell (GC) and arrow indicates basal GC. bar represents 10 µm. Right: Fluorescence intensity of 5mC plus 5hmC in central and basal germ cells from DEHP-BPA (DE/BP)-exposed testes or control (CTL) testes. Dotted lines represent median fluorescence intensity (CTL and DE/BP) in peritubular (orange) and sertoli cells (cyan).*** p ≤ 0.001 (Student’s t test_two-tailed). Violin Plot combined with box plot (median and interquartile ranges) from 10 to 180 cells per xenograft (from 6 individuals paired testes). **(B)** Volcano plot showing of differently methylated genomic regions (DMR) from DEHP/BPA (DE/BP) vs control (CTL) xenografted whole testes. Blue dots indicate hypermethylated DMRs (differential methylation >= 25, qValue ≤ 0.00005). Blue dots indicate hypomethylated DMRs (differential methylation ≤ 223 cl:1046 [Edit].25, qValue ≤ 0.00005). **(C)** Left and right plots: Violin plots showing the percentage of CpGs by genes (from promoter to 3’ end, left) or by gene regions (right plot) significantly differentially hypomethylated (hypom), hypermethylated (hyperm) or non modified (No Diff) in DE/BP testes compared to CTL. Upper numbers represent the percentage of genes presenting more than 50 % of CpG differentially hypomethylated or hypermethylated in DE/BP condition. **(D)** Mean expression of genes with > 50 % of hypomethylated CpG in promoter using ScRNAseq data from developing human gonads. **(E)** Functional enrichment analysis using Rare_Diseases_GeneRIF_Gene_Lists database in EnrichR of genes expressed by GCs (from undifferentiated PGCs to quiescent pre-spermatogonia) and showed >50% of hypomethylated CpG in promoter.

### DE/BP exposure causes defective spermatogenesis and seminoma formation

A number of hallmarks of seminomatous TGCTs were observed in germ cells from DE/BP testis: enrichment of GC with PGC features, GC DNA hypomethylation, fibrosis in the peritubular wall. As an illustration, we found that the transcriptomic profile observed previously in DE/BP testis (compared to CTL) correlates to the published transcriptomic profile of TGCTs (compared to control adult testis) with a global down-regulation of meiotic genes (Supplementary Figure 7). However, these TGCT features are primarily caused by differentiation defects in the GC and cannot be used to predict tumorigenicity. By stimulating spermatogenesis in DE/BP-treated GC, we would evaluate their differentiation ability. To accomplish this goal, human fetal GC was heterologously transplanted into adult mouse testis previously depleted of GC by busulfan treatment (Supplementary Figure 8A). In this model, hGC can relocate to the basal membrane of the murine seminiferous cords and generate and maintain colonies of spermatogonia. However, xenotransplanted hGC did not undergo complete spermatogenesis in murine testis with arrest at the meiotic stage (Liang et al., 2022; Nagano et al., 2002). To retrieve all GC populations in xenografted testis (*i.e*. pluripotent PGC, differentiating PGC and pre-spermatogonia), we cell-sorted β2-Microglobulin (B2M) negative cells at the end of exposure. The expression of B2M is low in germ cells compared to somatic cells, and GC are enriched in B2M-negative populations (Supplementary Figure 8B-D). B2M negative cells were transplantated into mouse adult testes chemically depleted of GC and, four weeks later, recipient testis was collected to performed histological analysis. As partial restoration of spermatogenesis could be observed 4 weeks after xenotransplantation (8 weeks after busulfan treatment) we identified human cells using an anti-human mitochondrial antibody. In the mouse seminiferous tubules, most human transplanted cells have germ cell features (from spermatogonia to pachytene spermatocytes) but few somatic cells as Sertoli cells could also be seen ( Figure 4A, Supplementary Figure 8E). DE/BP treatment seems to affect hGC survival and their ability to home at the plasma membrane as we observed significant increased in abnormal hGC (cell death, multinucleated cells or luminal cells) previously treated to DE/BP (Figure 4B). In addition, spermatogenesis progression seems also affected by previous DE/BP exposure as we observed an enrichment of undifferentiated cells in detriment to meiotic cells. First, we observed fewer hGC with meiotic features in the testis xenotransplanted with DE/BP treated cells than control (Figure 4C). Second, in xenotransplanted testis with DE/BP treated hGC, we found a significant increase of the number in PLAP expressing human cells (Figure 4D). Placental alkaline phosphatase (PLAP) is one of the cellular phosphatases expressed in seminoma and carcinoma. PLAP is also expressed by human fetal GC with a decrease expression along GC differentiation. There are a few PLAP-positive GCs that can be retrieved from neonatal testis after birth, but these cells disappear after a few months (Li et al., 2023). In some cases, we could observed malignant neoplasms in testes after hGC transplantation (Figure 4E-F, Figure 5). Besides testis and epididymis, human cells are also found invading the retroperitoneum outside of the testis. The invasion of human cells was extremely rapid in these cases, and we were able to observe scrotal skin lesions in 4 weeks following the transplant (Supplementary Figure 9E). There is an expansion of intra- and extra-tubular human cells in the testis, causing the seminiferous epithelium to disorganize. Loss of vimentin expression by Sertoli cells was observed in these disrupted seminiferous tubules (Figure 5A). As a result of hypoxia and global inflammation in the testis, we detected homogeneous human cell death in the testis, as illustrated by cleaved caspase 3 staining presented in Figure 5A and presence of pyknotic cells inside seminiferous tubules (Figure 4F-G). However, human cells outside the testis that remain alive presented some typical features of seminoma. (Figure 4F). These cells were homogeneous, round and had vesicular or polygonal nuclei with prominent nucleoli. They expressed seminoma markers and meiotic markers such as *PLAP, BRAP, ASB9, SYCP3, HORMAD and DAZL* and remain proliferative as illustrated by PCNA staining (Figure 5B). Altogether, these results indicate the development of seminoma and the incidence of seminoma formation was increased with DE/BP treated hGC in comparison ti CTL treated hGC (Figure 5E, 8 cases on 22 transplantations for DE/BP cells versus 2 cases on 22 transplantations for CTL cells). These findings demonstrated that DE/BP exposure could permanently damage fetal GC disrupting their ability to initiate spermatogenesis and promoting their tumorigenesis.

**Figure 4.**
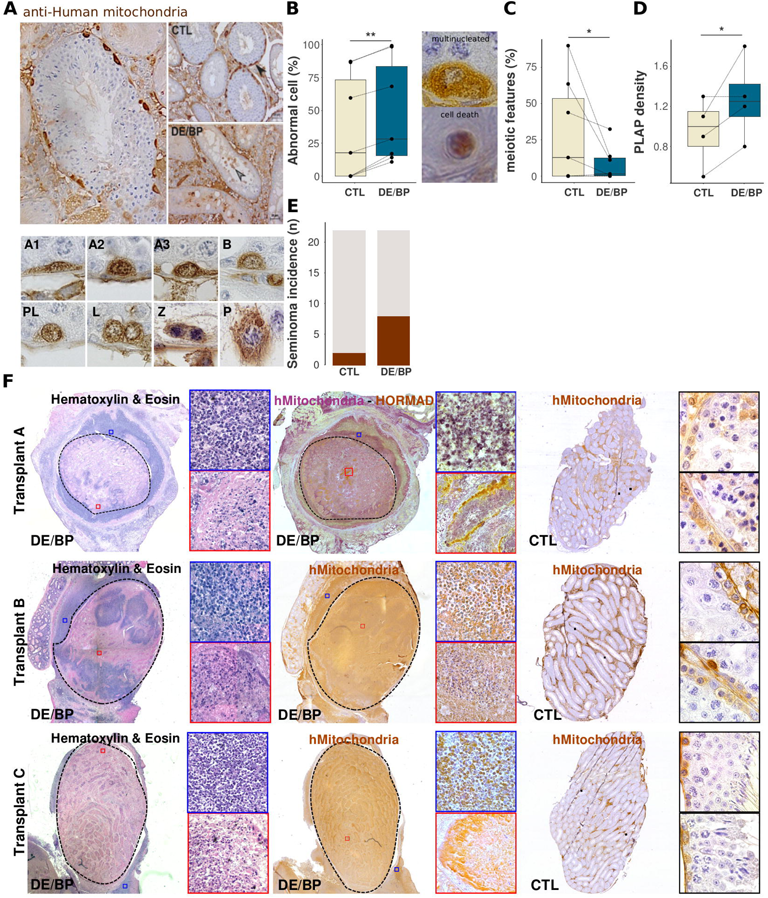
Spermatogenesis defects and malignant transformation with DE/BP treated GC. **(A)** Representative images of testes 4 weeks after human GC (hGC) xenotransplantation. HGC were detected by immunostaining with anti-human mitochondria antibody (brown). Lower panel: representative image of hGC toward spermatogonial differentiation according to the nucleus shape and chromatin condensation as described by Clermond Y. A1-3: Type A dark (A1-2) and pale (A3) spermatogonia, B: Type B spermatogonia, PL: Preleptonema, L: Leptonema, Z: Zygonema, P: Pachynema (Clermont, 1966). **(B)** Percentage of human abnormal cell (/total human cells) in seminiferous tubules of murine testis xenotransplanted with CTL and DE/BP human B2M negative cells. Multinucleated cells, cells with pyknotic nucleus and/or breakdown of the plasma membrane or luminal cells are considered as abnormal cells. **(C)** Percentage of basal human cells harboring meiotic features (/ total basal cells). **(D)** Number of PLAP positive cells per seminiferous tubules. **(B-D)** box plot (median and interquartile ranges) from 4-6 individuals paired xenotransplanted testes. * p ≤ 0.05, ** p≤ 0.01, ns: non significant (Paired student’s t test). Grey dotted line connects paired testis transplanted with cells from the same embryo (indicated by points). **(E)** Incidence of seminoma formation after transplantation of murine testis with CTL- and DE/BP-treated B2M negative cells. (**F**) Representative pictures of three seminoma formation or normal development after DE/BP or CTL human GC transplantation (Transplant A-C: “paired” seminoma/normal testis obtained by DE/BP or CTL hGC transplantation from the same human testis). Left panel: haematoxylin-eosin staining in seminoma induced by DE/BP hGC transplantation. Middle panel: Immunostaining of anti-human mitochondria in seminoma induced by DE/BP hGC transplantation. Right panel: Immunostaining of anti-human mitochondria in normal testis after CTL hGC transplantation. For each panels: left magnifications of cells inside (red square) or outside (blue square) testis.

**Figure 5:**
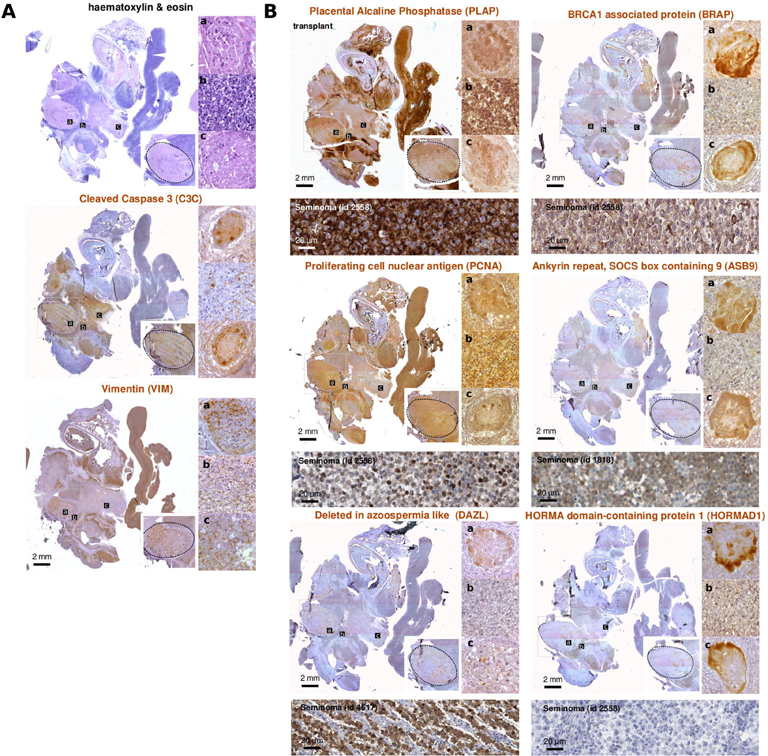
Illustration of seminoma/proliferative and apoptotic profiles in hGC in a testis presenting an hight malignant transformation and extratesticular cell invasion. **(A)** H/E staining, cleaved-Caspase3 (C3C) and Vimentin Immunostainings (VIM) **(B)** Immunostainings of undifferentiated fetal GC (PLAP) and proliferative cells (PCNA), meiotic cells (DAZL, HORMAD) and TGCT and seminoma markers (ASB9, BRAP). Upper panel: Immunostaining of seminoma extracted from Human protein Atlas (same references of antibodies were used for characterization of seminomatous formation in xenotranplanted testis, the id correspond to the human protein atlas id).

## DISCUSSION

Parallel temporal trends in testicular cancer and other male reproductive disorders led to the hypothesis of a common environmental cause for several diseases of the male reproductive system (Juul et al., 2014; Skakkebaek et al., 2016). In 2001, Testicular Dysgenesis Syndrome (TDS) was first described encompassing poor semen quality, undescended testis, hypospadia and testicular germ cell cancer (TGCT) (Skakkebaek et al., 2001). TDS was the hypothesized result of disrupted embryonal programming and gonadal development by environmental and/or genetic causes (Juul et al., 2014). If numerous experimental and epidemiological models evidenced the impact of gestational EDCs exposure on steroidogenesis, androgen response and masculinization, the impact of gestational EDCs exposure on TGCT incidence was unclear (Fénichel et al., 2018; Kilcoyne and Mitchell, 2019; Sweeney et al., 2015; van den Driesche et al., 2017). However, despite a clear association between subfertility and increased risk of TGCT, epidemiologic studies provide conflicting support for the existence of a link between EDCs exposure, germ cell failure during development and TGCT incidence (Akre and Richiardi, 2009; Faja et al., 2022; Fénichel et al., 2018; Rodprasert et al., 2021). Currently there are no experimental models to study the effects of EDCs on TGCT, nor are there rodent models that develop TGCT spontaneously. Asynchronicity of germ cell differentiation and long development time could explain human susceptibility to develop TGCT. In the present study, we propose an original experimental model that could link EDCs exposure during human gonadal development and increase risk to induce germ cell carcinogenesis. This could lead to seminoma formation.

We decided to use a mixture of the most common environmental EDCs: DEHP and BPA. In spite of recent restrictions on DEHP and BPA use in the European Union, we were always able to detect DEHP and BPA in serum and urine samples.. DEHP is retrieved in fetal cord blood samples at 1.1 × 10^−8^ M (Lin et al., 2011) and BPA concentration in maternal serum could be around 10^−9^ to 10^−12^ (Teeguarden et al., 2016). To mimic environmental exposure to EDCs, prolonged treatment with a mixture of DEHP and BPA (DE/BP, 1µM each) *via* xenograft model of human fetal testis was realized. According to the same protocol of exposure, plasma levels of BPA were 0.1 µM corresponding to environmental exposure (Eladak et al., 2018). In addition, DE/ BP exposure during several rounds of PGC to pre-spermatogonia transition (from 8 to 18 WPF) mimics *in utero* chronic exposure to this compounds. Using this model, we showed, by transciptomic and histological analyses, that DE/BP exposure affects PGC to pre-spermatogonia differentiation. DE/BP exposure led to enrichment of pluripotent PGCs, impairment of pre-spermatogonia differentiation, proliferation maintenance in pre-spermatogonia and precocious meiosis initiation. In addition, we observed global DNA hypomethylation in DE/BP treated GC. Interestingly, hypomethylation was observed in PGCs and in pre-spermatogonia suggesting that this event was not correlated to GC differentiation and could be imputable to DE/BP exposure. Interestingly, differentially expressed or methylated genes after DE/BP exposure are linked to TGCTs. Heterologous transplantation of DE/BP treated GCs leads to spermatogenesis defect and in some cases, seminoma formation. This is the first experimental evidence that EDCs exposure during fetal life could induce permanent defect in GC leading to differentiation impairment and seminoma formation. In addition to GC differentiation failure, we also observed that DE/BP exposure alters somatic cells in the fetal testis. Indeed, a transcriptional signature of fibrosis and peritubular wall thickness was observed in DE/BP testis. This suggests a modulation of TGFb signaling in these cells and this was confirmed by ScRNAseq analysis (Chung et al., 2021).

Unfortunately, we did not identify molecular mechanisms involved after DE/BP exposure leading to germ cell differentiation defects and seminoma formation. As fetal GC differentiation depends on intrinsic/epigenetic and somatic signaling we could not exclude the impact of DE/BP induced somatic defects on GC differentiation (Frydman et al., 2017; Hargan-Calvopina et al., 2016; Hargy and Sasaki, 2023; Webster et al., 2021). Regulation of fetal male germ cell development is highly dependent on TGFb signaling. Members of this superfamily are directly involved in a range of cellular processes including maintenance of pluripotency and prevention of meiosis initiation and mitotic arrest (Moreno et al., 2010; Souquet et al., 2012; Spiller et al., 2017). Environmental stress responsible for modulating TGFb signaling may also alter TGFb signaling in GC, affecting differentiation. We also proposed that DE/BP-induced somatic alterations illustrated by a decrease in PMC proliferation could prevent somatic production of factors that prevent GC male pathway. Interestingly, peritubular wall thickness and GC alteration was similarly observed after gestational exposure to dibutylphthalate in rats but no mechanism explaining this phenotype was proposed by the authors (Fisher et al., 2003). This observation suggests common mechanisms between EDC compounds causing fibrosis, testis cord disorganization leading to impairment of the stem cell niche and somatic production of paracrine factors involved in GC differentiation. Impairment of GC differentiation could also be due to modification directly in GC. According to previous research, BPA analogues can impair GC differentiation in ovary by causing oxidative DNA lesions (Abdallah et al., 2023). Oxidative lesions such as 8-oxodeoxyguanine (8-OdG) near CpG islands inhibits the DNMT binding and could induce TET1 activity leading to passive or active demethylation (Hahm et al., 2022). Modification of the DNA methylation landscape, as well as, the presence of 8-OdG in promoter regions could also have consequences for the set-up of the transcriptional differentiation program in GC. Altogether, oxidative stress induced by DE/BP exposure could explain the fibrotic signature, GC DNA hypomethylation and failure of GC differentiation. Further studies would confirm the pro-oxidant impact of DE/BP and correlate germ cell failure with DNA oxidative lesions. However, one of the intriguing questions is whether exposure of PGCs to stress conditions would induce carcinogenesis and seminoma formation. Teratomas are well known to result from nondifferentiation of pluripotent PGC. It was observed after genetic invalidation of Dnd1, Nanos2 or Dazl in mice (Chen et al., 2014; Imai et al., 2020; Webster et al., 2021) and in the case of allogeneic transplantation of embryonic stem cells or migrating PGC in post-natal testis form teratomas (Clark et al., 2017; Hentze et al., 2009). However, we could not observe teratoma formation after germ cell transplantation even with DE/BP than CTL treated cells. This is despite the presence of pluripotent PGC in xenografted testis. For this reason, we could speculate that malignant transformation could originate from abnormal or proliferative pre-spermatogonia. Moreover, malignant transformation and seminoma formation were not observed systematically. This suggests that DE/BP exposure could induce stochastic modifications in GC resulting in their carcinogenesis and tumorigenesis. Numerous genome-wide association studies and meta-analysis have identified TGCT susceptibility loci. In this study, we did not find transcriptional and epigenetic modifications in these loci (Supplementary Figure 9). This excludes a DE/BP induced genomic modification preferentially in one of these loci explaining seminoma formation (Pluta et al., 2021). TGCTs and seminoma are consistently aneuploid and in the testis, the seminomas have a mean hyper-triploid DNA index (Shen et al., 2018). Aneuploidy and unrepaired DNA strand breaks could cause cytokinesis failures exacerbating chromosome instability and aneuploidy (Lens and Medema, 2019). In DE/BP-treated GCs transplanted into murine testis we observed an increase in cytokinesis failures and cell death. It is possible that these defects were originally caused by genome duplications and mutations that promote tumorigenesis and seminoma formation. To link with previous comment, 8-OdG formation in the genome is highly mutagenic and leads to genome instability and cancer (Hahm et al., 2022).The genetic and epigenetic instability caused by DE/BP-induced DNA oxidative damage during fetal differentiation may explain mutagenesis later in life. Unfortunately, due to low cell number or low DNA quality due to GC death inside the testis and in the case of seminomatous growth, no genome sequencing in the pre-tumorigenesis and seminomatous cells have been conducted to identify mutation hot-spots, whole genome duplication and clonal heterogeneity. It will be critical to evidence genomic modification (i.e. DNA oxidation, DNA mutation) in fetal GCs just after DE/BP exposure to attribute these modifications to seminoma formation.

In spite of the lack of molecular mechanisms induced by DE/BP exposure, the present study is the first to demonstrate that EDC exposure during human fetal testis development results in tumorigenesis and TGCT.

## Acknowledgments

We are grateful to A. Gouret and A. Leliard for her skillful secretarial assistance. We also thank the team within the animal housing facility at the iRCM. This research was supported by the Plan Cancer Inserm (INSERM) and Université de Paris and INSERM. A.J. were supported by a fellowship from the Ministère de l’Enseignement et Recherche.

## SUPPLEMENTARY FIGURE LEGENDS

**Supplementary Figure 1: *In vivo* germ cell differentiation during development.** *In vivo* testes from 8 to 35 WPF were collected and germ cell (GC) differentiation was assayed. **(A)** percentage of AP2γ and OCT-3/4 positive GC during development. **(B)** percentage of DDX4 and MAGEA4 positive GC during development. **(C)** ratio of AP2γ/MAGEA4 positive GC during development. Each dot correspond to one testis. For A,B and C, green and orange dots correspond to 8-11 or 25-34 WPF testes used in Figure 1.

**Supplementary Figure 2. Maintenance of germ cell pluripotency after DE/BP exposure. (A)** Representative images of testes stained with anti -PDPN (green) and -KIT (red) antibodies counterstained with DAPI (blue). Bar represents 10 µm. **(B)** Percentage of KIT- and PDPN-positive GCs (/total GCs) from CTL and DE/BP xenografted testes. Box plot (median and interquartile ranges) from 6-8 individuals paired xenografted testes. * p ≤ 0.05 (paired student’s t test). Grey dotted line connects paired xenografted testes (indicated by points).

**Supplementary Figure 3. scRNAseq analysis in DE/BP and CTL testes. (A)** Two-dimensional UMAP (Uniform Manifold Approximation and Projection) plot of testicular cells from 2 paired testes treated to DE/BP or solvant (CTL_A, DE/BP_A and CTL_B, DE/BP_B). Cell clusters are colored according to the cell identity. **(B)** Number of retrieved cells in condition and cluster after encapsulation and cell filtering. **(C)** UMAP of somatic cells (supporting and intertistial cells) after downsampling in CTL condition. **(D)** Expression of markers of sertoli (*SOX9*) and interstitial (*NR2F2*, *INSL3*) cells. **(E)** VolcanoPlot of differentially expressed genes in DE/BP treated cells compared to CTL cells. Red and green dots indicate genes involved in pathways enriched after functional analysis by EnrichR. Red dots indicate genes involved ECM and focal adhesion pathways. Green dots indicates genes involved in ECM regulation by TGFβ signaling.

**Supplementary Figure 4. Use of published scRNAseq data from human developing gonad for bulk RNAseq deconvolution. (A)** Violin Plot presenting gene expression of specific markers of pluripotent and differentiating PGCs, meiotic oocytes, quiescent pre-spermatogonia, supporting cells (sertoli cells), interstitial cells (peritubular and leydig cells), and immune cells (myeloid, lymphoid and endothelial cells), **(B)** distribution of cell density in all cell clusters expressing high, medium and low levels of down-regulated (upper, in blue) or up-regulated (lower, in red) genesets after DE/BP exposure observed in bulk RNA-seq analysis. **(C)** Heatmaps showing differential expression between DE/BP condition vs CTL condition of genes expressed by germ and somatic cells.

**Supplementary Figure 5. DE/BP exposure affects peritubular cell proliferation. (A)** Representative images of testes stained with anti-PCNA (green) antibody and counterstained with DAPI (blue). Arrowhead indicates proliferative peritubular cells. In left small panels: higher magnification with circled peritubular cells. bar represents 10 µm. **(B)** Percentage of PCNA positive peritubular cells.* p ≤ 0.05 (Wilcoxon’s t test_paired). **(C)** Number of peritubular cell layers per peritubular wall (dot represents mean of 30 quantification per testis). ns: non significative (Wilcoxon’s t test_paired, pval= 0.1094). For (B) and (C). Bar Plot (median and interquartile ranges from 7 individual paired testes) combined with dot plot. Each linked dots represent paired testes (CTL and DE/BP).

**Supplementary Figure 6. Analysis of DNA methylation by immunostaining of 5mC and 5hmC in testicular cells in DE/BP and CTL testes. (A)** Graphical representation of germ cell migration and differentiation during fetal development. Pluripotent and differentiating PGCs are mostly located in the middle of the testis cords. During differentiation, positive pre-spermatogonia migrates to the basal compartment and express MAGE-A4. **(B)** Fluorescence intensity of 5mC and 5hmC in germ (central and basal germ cells) and somatic cells (peritubular, sertoli and leydig cells) of DE/BP and CTL testes. No significant difference between treatment was observed by Student’s t test_two-tailed. **(C)** Fluorescence intensity of 5mC and 5hmC in central and basal germ cells from DEHP-BPA (DE/BP)-exposed xenografted testes or control (CTL) testes in function of the initial age of embryo (from 7.7 to 10.9 WPF [week post-fertilization]). Violin Plot combined with box plot (median and interquartile ranges) from 10 to 180 cells per xenograft.

**Supplementary Figure 7. Co-profiling of DEGs observed after DE/BP exposure and in testicular germ cell tumors (TGCTs).** LogFCs of DEGs in DE/BP xenografts (compared to CTL xenografts) were compared to LogFC of DEGs in TGCTs (compared to normal tissue, data obtained from Chang et al study (Chang et al., 2019). Colored circles correspond to DEGs with qvalue <=0.05 in both transcriptome analyses. Dot size: −log10(sum of qvalues). Red circles highlight some of genes normally involved during meiosis such as *SYCP3*, *SYCP1*, *RAD51*, *SMC1B*, *RBM44*, *PIWIL*, *SPATA22*, *HORMAD1*, *MSH4*.

**Supplementary Figure 8. Xenotransplantation of human cells from xenografted testis in murice testis recipient. (A)** Schematic representation of the experimental design of xenotransplantation of human cells in murine testis. Human fetal testis was grafted in the back of 2 nude mice. One weeks later, xenografted mice were exposed to DEHP-BPA (DE/BP) mixture or to vehicle during 7-10 weeks (depending of the age of the human fetus). At the end of exposure, human testes were collected, dissociated and β2 microglobuline (B2M) negative cells (B2M neg, mostly germ cells) were sorted by FACS and then injected via efferent duct into the seminiferous tubules of the testis of two recipient mice (one by condition). To deplete endogenous spermatogenesis, the recipient mouse have previously been treated to bulsulfan. Four weeks after xenotransplantation, testes were collected for histological analyses. **(B)** Expression of *B2M*, *POU5F1*, *KIT*, *DDX4* and *MAGEA4* in germ and somatic cell cluster from published scRNAseq data. uGC: undifferentiated germ cells, dGC: differentiating germ cells, qGC: quiescent germ cells, Sup: supporting somatic cells (Sertoli cells), Int: Interstitial somatic cells (Leydig and Peritubular cells), Imm:Immune cells (lymphoid, myeloid and endothelial cells). **(C)** Gating strategy of B2M negative cells. Only viable cells were conserved using HO 33358. **(D)** Relative quantification (RQ) of gene expression of germ (left) and somatic (right) cells in B2M negative cell population compared to the B2M positive cell population. **(E)** Representative images of testes 4 weeks after xenotransplantation immunostained with anti-human mitochondria antibody (brown) to detect human cells. a: unmagnified image of testis section. From b to d: representative images of spermatogonia-like germ cells (b), spermatocyte-like germ cells (c), luminal cells (d) and sertoli-like cells (d). Bar represents 15 µm. **(F)** Upper: Picture of skin lesion (rectangle) due to metastasized germ cell expansion. Lower: Picture of urogenital organs four weeks after xenotransplantation. Note the irregular shape of testes encircled by dotted lines. Arrowhead indicates local cell extravasation outside the testis. bl: bladder, te: testis, ep: epididymis.

**Supplementary Figure 9. Differential expression and DNA methylation of susceptibility genes associated with testicular germ cell tumors. (A)** LogFCs in DE/BP xenografts (compared to CTL xenografts) of target genes moderately or highly associated to testicular germ cell tumor (TGCT) as proposed by the testicular cancer consortium. Dot size: −log10(qvalues). Blue dot: significant downregulation (LogFC ≤ −0.5, qValue ≤ 0.05). Red dot: significant upregulation (LogFC ≥ 0.5, qValue ≤ 0.05). Significant differentially expressed genes (26.9 % of all genes associated to TGCT) are indicated below (genes highly associated to TGCT) or above (genes moderately associated to TGCT) dots. Same text colors (green, blue, pink, purple and orange) indicates genes located in the same chromosomal region (in distance of 10 Kb)/ Random sampling: 23.9± 4.4 % random genes (10^3^ sampling of 90 genes) were significantly differentially expressed. **(B)** Percentage of CpGs by genes (from promoter to 3’ end, left) differentially significantly hypomethylated (negative percentage) or hypermethylated (positive percentage). Dot size correspond to the number of CpGs sequenced by genes. Differentially methylated genes (with more than 30 % of significant hypo- and/or hyper-methylated CpG, 16.3 % of all genes associated to TGCT) are indicated below (genes highly associated to TGCT) or above (genes moderately associated to TGCT) dots. Same text colors indicates genes located in the same chromosomal region (in distance of 10 Kb). Random sampling: 20.9± 4.2 % random genes (10^3^ sampling of 90 genes) were significantly differentially methylated.

