## Supplementary figures and images for "Evidence of the ability of endocrine disrupting compounds to induce testicular germ cell cancer in humans"

### Supplementary Figure 1

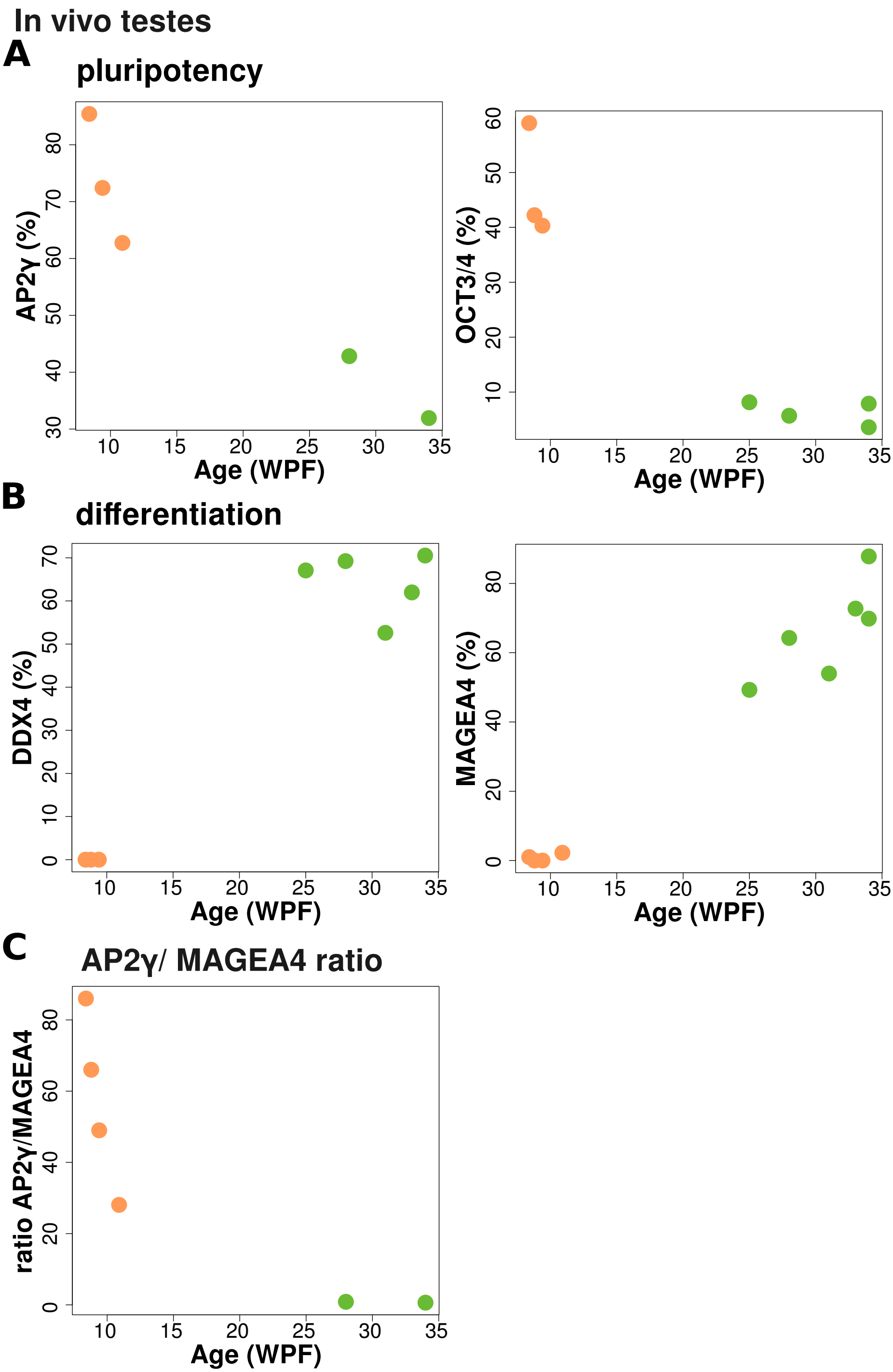

### Supplementary Figure 2

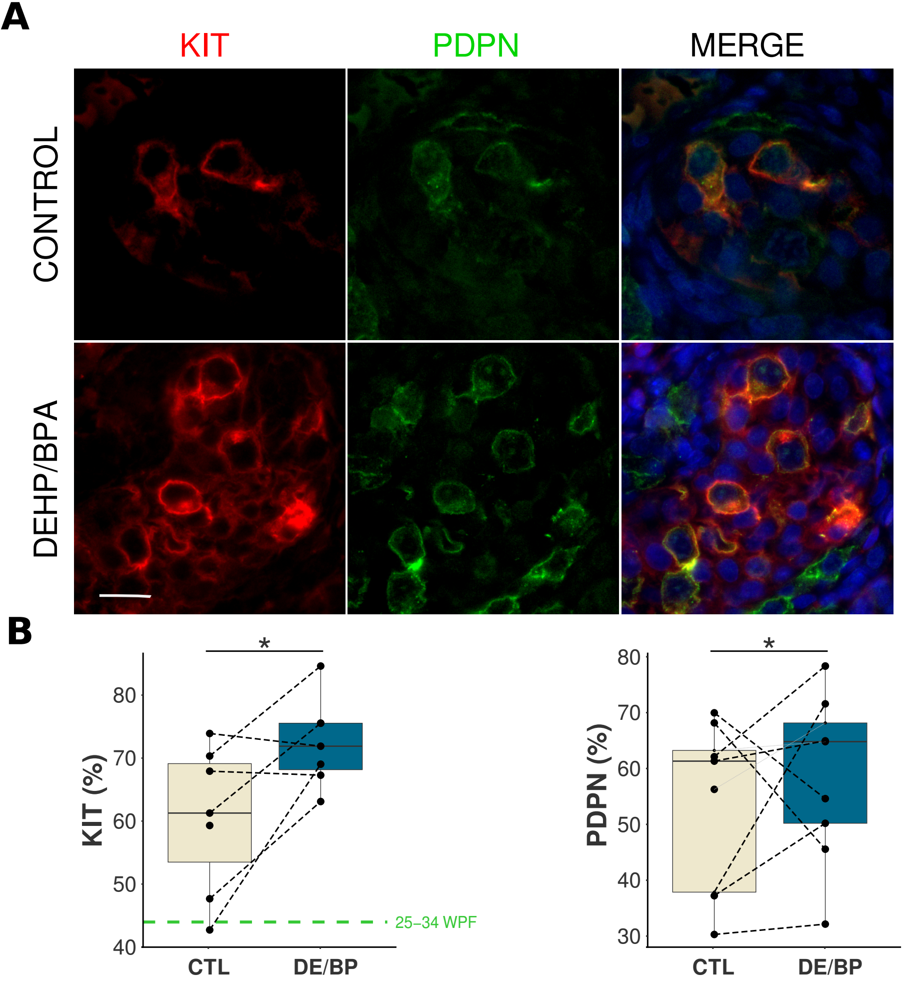

### Supplementary Figure 4

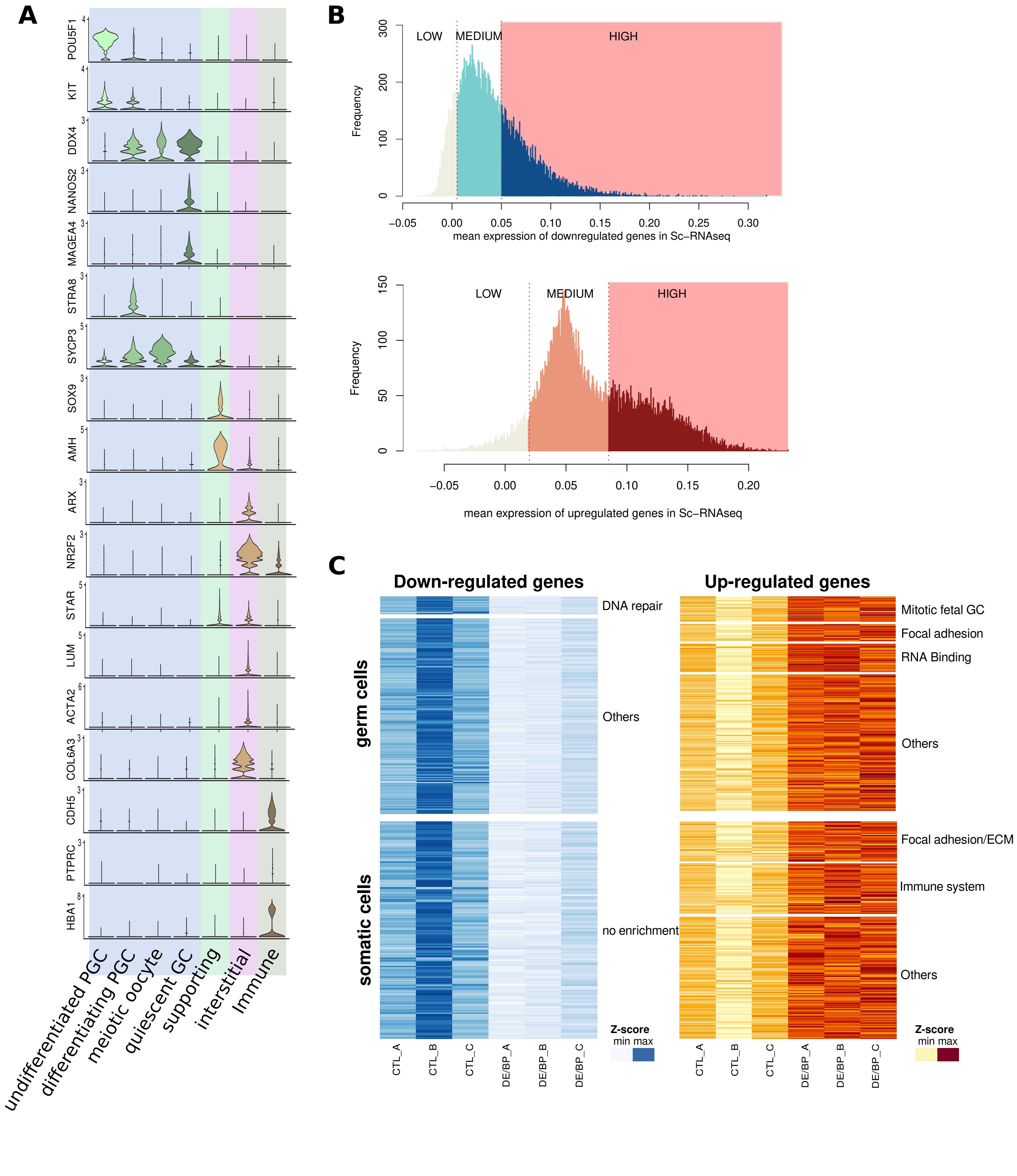

### Supplementary Figure 5

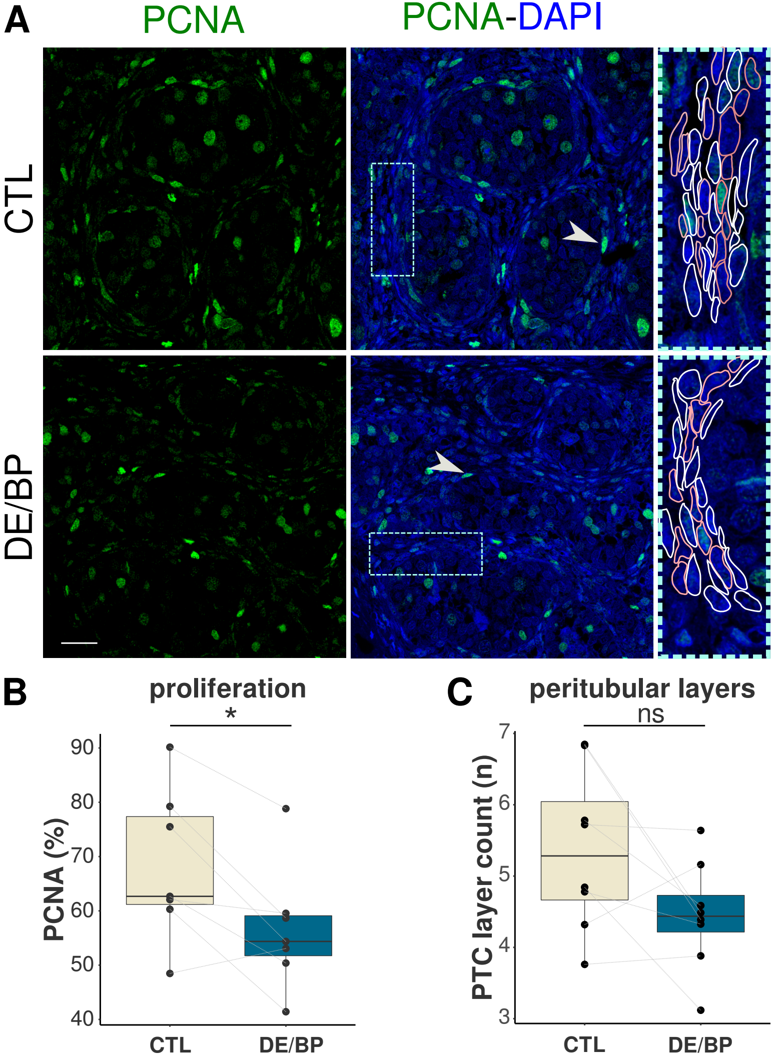

### Supplementary Figure 6

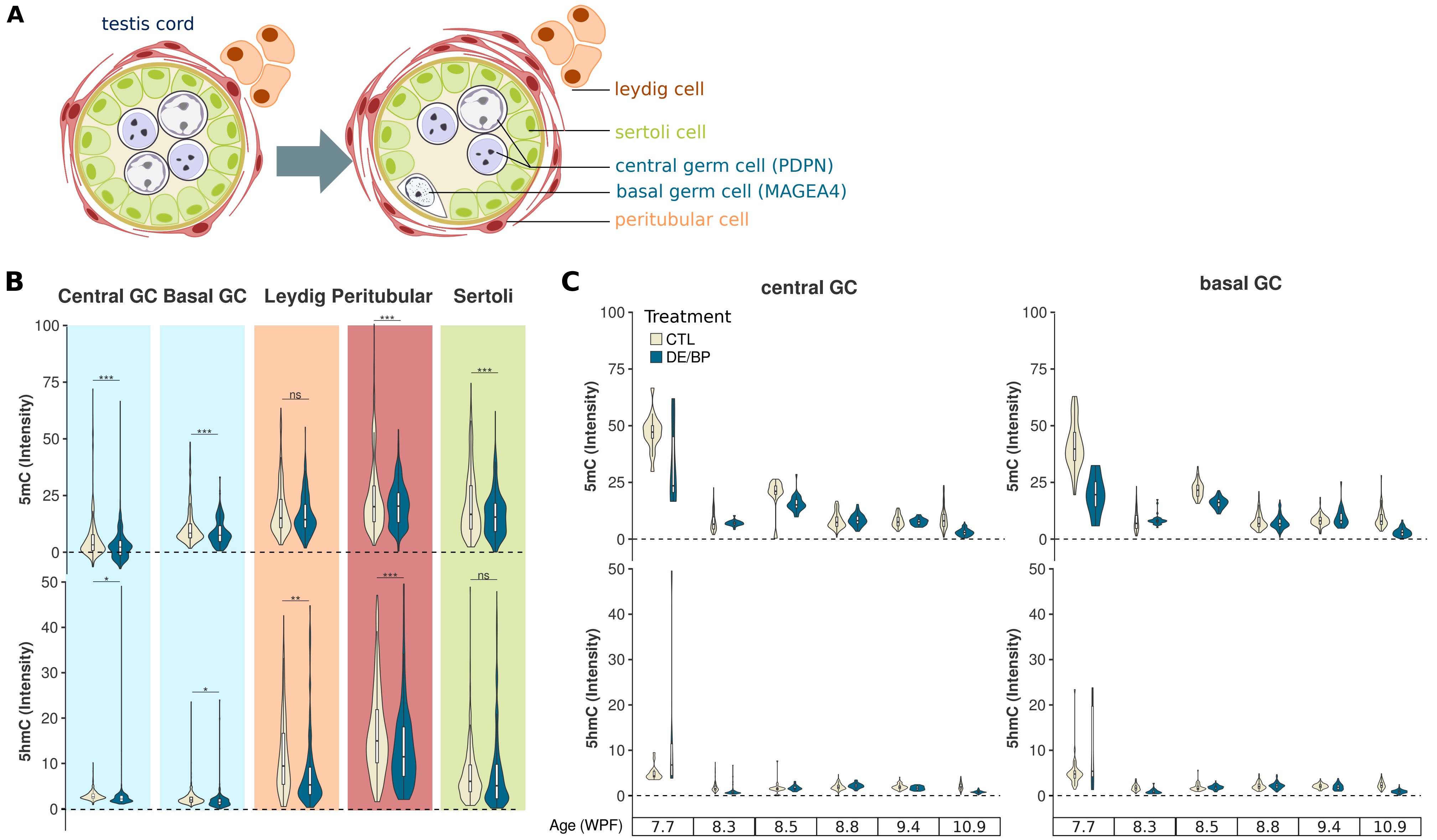

### Supplementary Figure 7

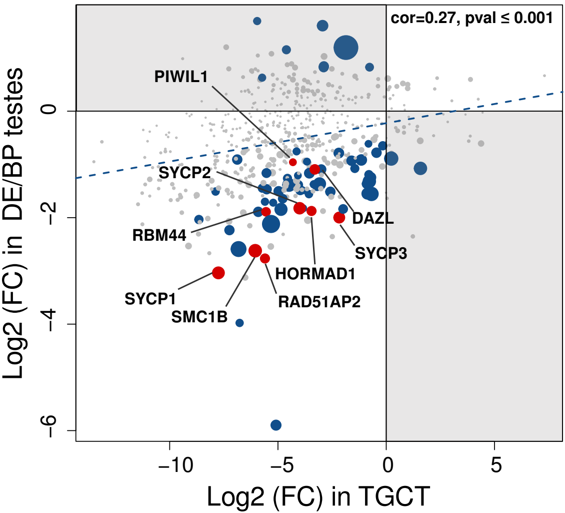


Chang Y. et al. Cancer Med. 2019

### Supplementary Figure 8

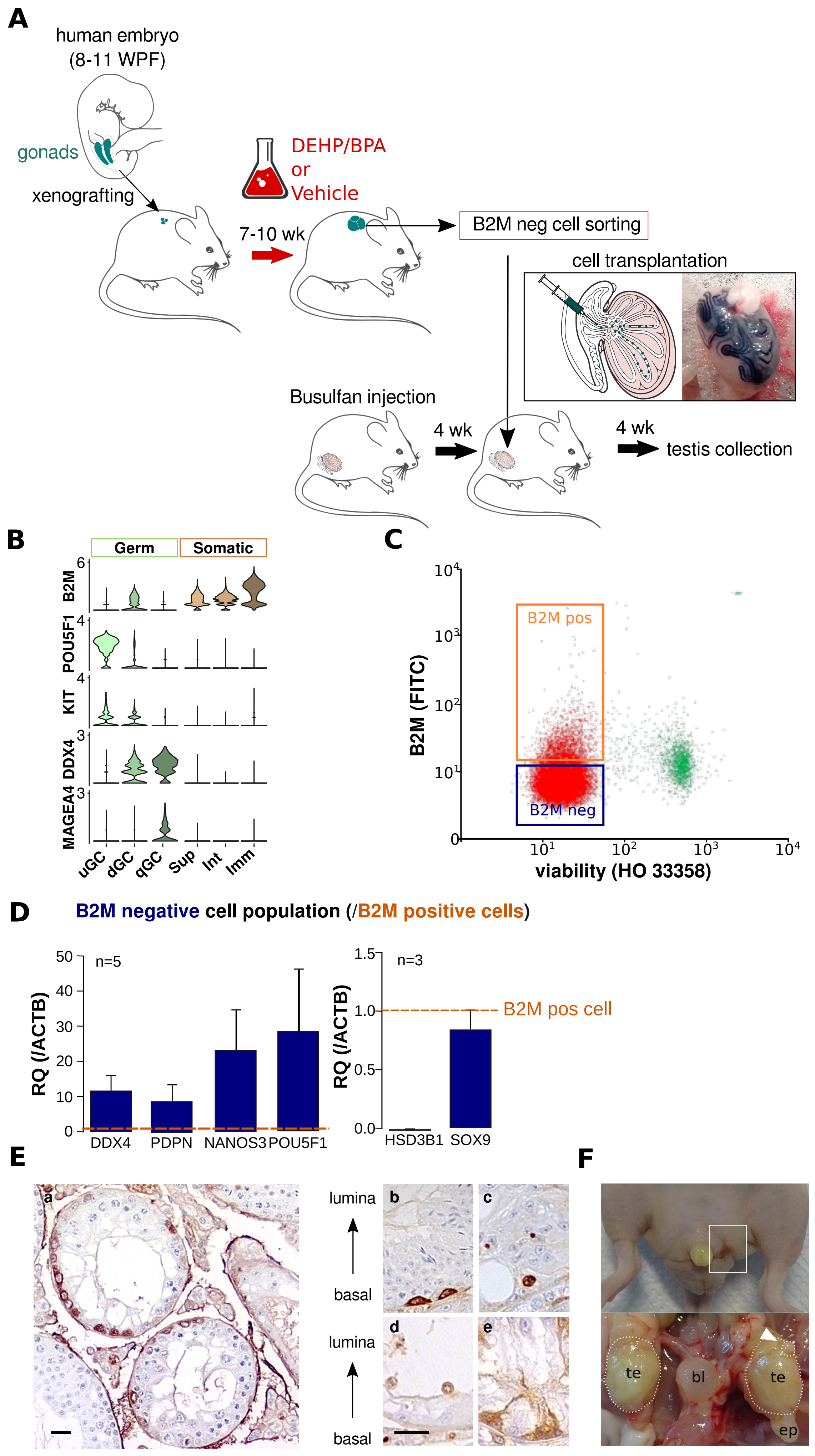

### Supplementary Figure 9

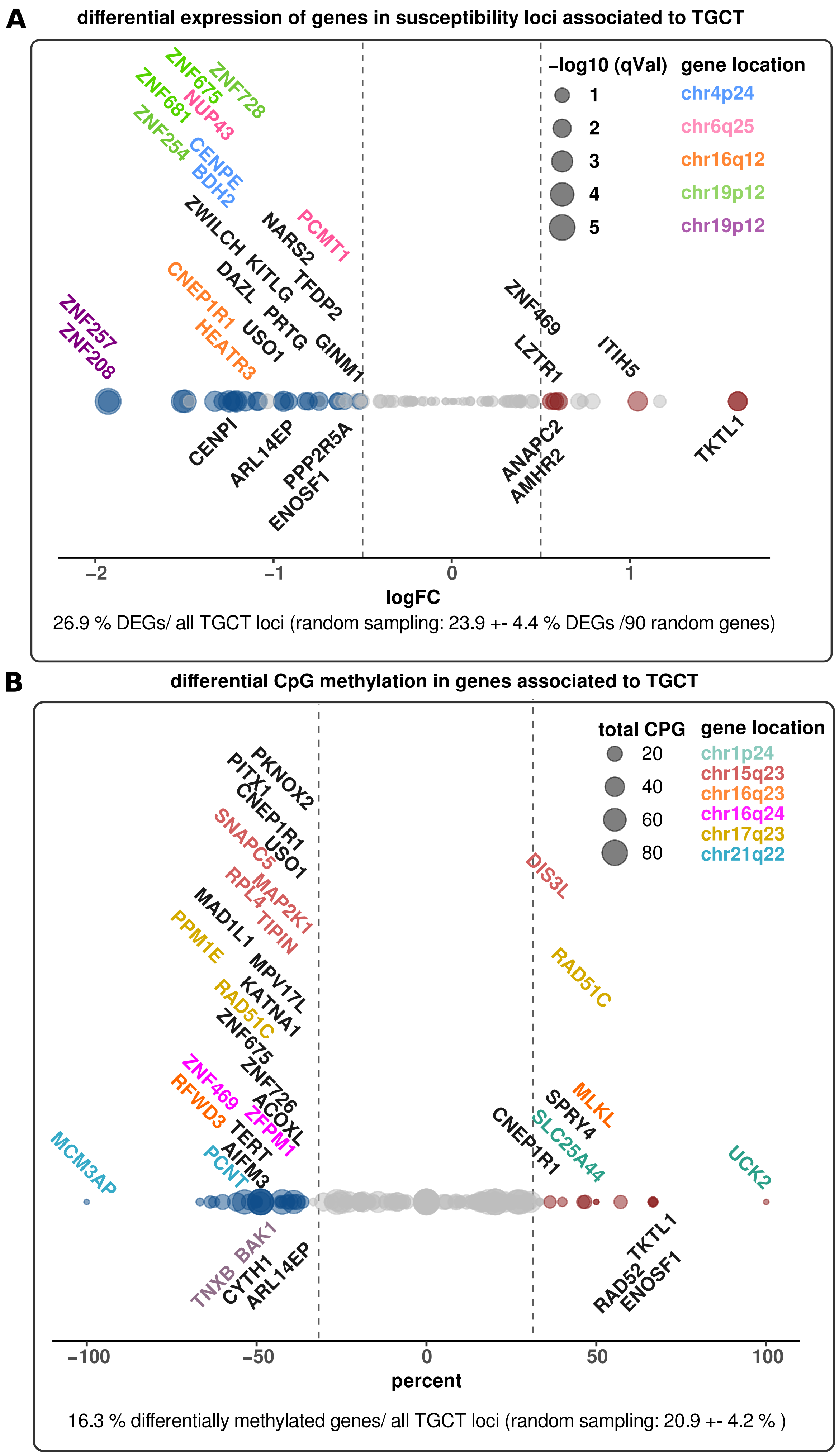
