## Supplementary Figure 3 for "Evidence of the ability of endocrine disrupting compounds to induce testicular germ cell cancer in humans"

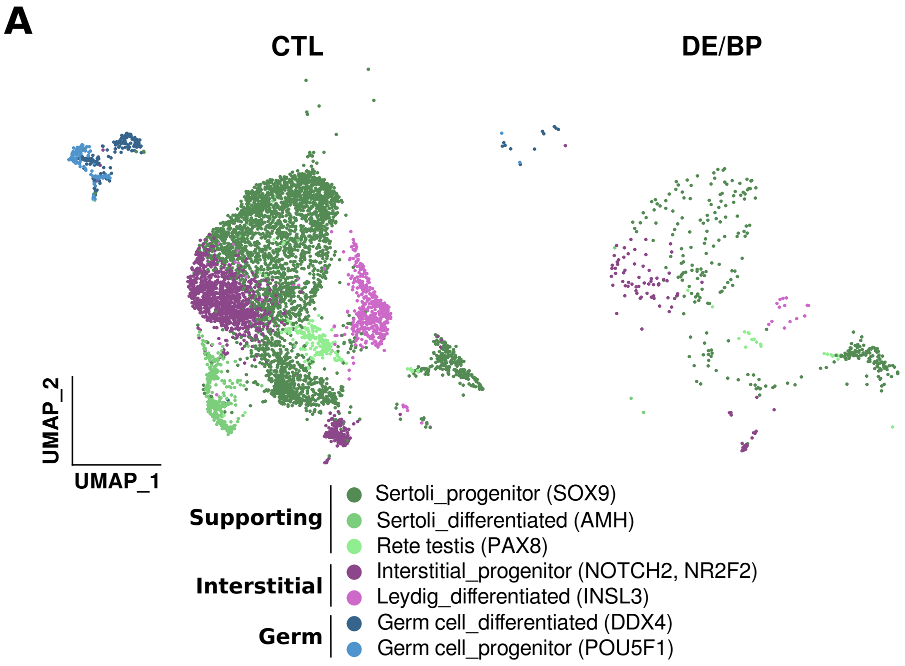


**B**

| | CTL_A | DE/BP_A | CTL_B | DE/BP_B |
|---|---|---|---|---|
| Sertoli_progenitor | 1024 | 69 | 2333 | 230 |
| Sertoli_differentiated | 63 | 2 | 260 | 1 |
| Rete testis | 12 | 1 | 217 | 20 |
| Interstitial_progenitor | 284 | 46 | 1036 | 50 |
| Leydig_differentiated | 225 | 6 | 251 | 8 |
| Germ cell_differentiated | 35 | 6 | 125 | 5 |
| Germ cell_progenitor | 46 | 0 | 89 | 2 |

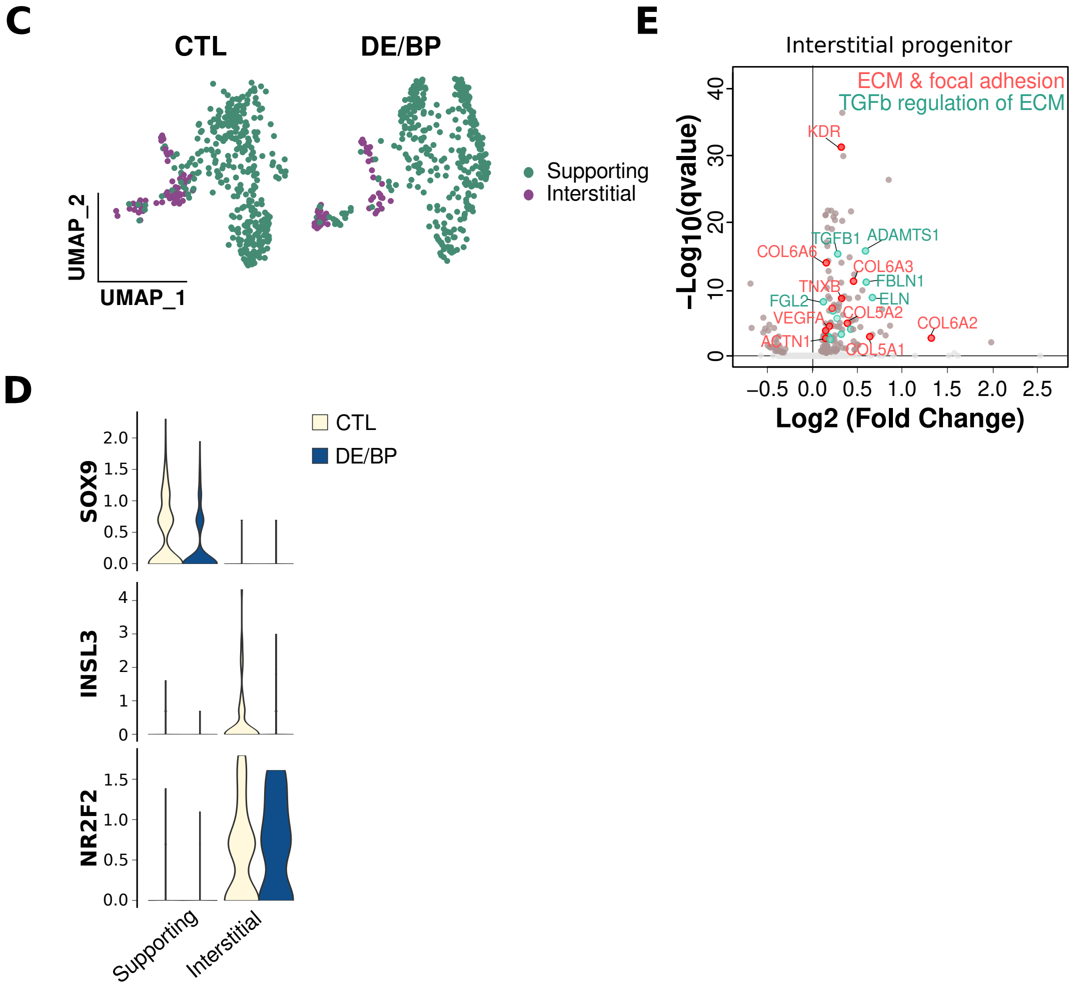
